# Decoupling Choice from Motor Response Reduces Choice-History Effects

**DOI:** 10.64898/2026.07.02.736214

**Authors:** Sriram Thothathri, Bharath Chandra Talluri, S. Shushruth, Michael N. Shadlen, Hendrikje Nienborg

## Abstract

Perception and action form a loop as an organism interacts with its environment. Here, we probe the interdependence between the two in the context of choice-history effects, the tendency of past choices and their outcomes to bias current choices. We investigated if the modulation by previous choices depended on the coupling between perceptual choices and the motor responses reporting these choices. We analyzed data from non-human primates (*Macaca mulatta*; three females and five males) performing two different perceptual decision-making tasks. Each task had a “coupled” variant with a fixed mapping between perceptual choices and motor plans used to report the choice, and an “uncoupled” variant, in which this mapping varied randomly across trials. We found that, in both tasks, the animals in the coupled variant had larger choice-history effects compared to the animals in the uncoupled variant. Decoupling choices from motor responses was further associated with an inability by the animals to learn experimentally induced stimulus sequence regularities. Together, these findings identify choice-response coupling as a factor shaping animals’ ability to use recent history to guide behavior and highlight the tight link between cognitive and motor processes.

## Introduction

The ability of the brain to integrate prior experiences, including past decisions and their outcomes, into decision-making is fundamental to adapting to an ever-evolving, often ambiguous, environment. The resulting heuristics are thought to be beneficial in natural environments (Gardner, 2019) and persist in laboratory tasks aimed at identifying the neural and computational mechanisms of cognition (Rahnev and Denison, 2018). In perceptual tasks, perceptual decisions and their discriminanda are typically designed to be statistically independent of the preceding decisions. Yet, despite the designed statistical independence over time, the influence of previous decisions and outcomes on decision behavior has been identified in numerous studies across human and non-human species (e.g. Akrami et al., 2018; Busse et al., 2011; Fernberger, 1920; Fischer and Whitney, 2014; Fritsche et al., 2017; Mochol et al., 2021; Van Bergen and Jehee, 2019).

Studies have largely linked history effects to previous choices, previous stimuli, previous rewards or a combination of these (Akaishi et al., 2014; Kalhan et al., 2026; Pape and Siegel, 2016; Zhang and Alais, 2020), and found that human and non-human animals exhibit idiosyncratic biases by repeating or alternating previous choices or rewarded stimulus categories. However, many studies treat choices and the motor responses used to report them as equivalent, by having a consistent mapping between stimulus categories and the motor response. For example, in classical direction discrimination tasks, a leftward saccade is typically required to report leftward moving dots and a rightward saccade for rightward moving dots. The contributions of previous choices and previous motor responses to history effects can be decoupled with tasks that randomize the mapping between stimulus categories and motor responses across trials. Previous studies in humans reported variable findings using such tasks (Braun et al., 2018; Zhang and Alais, 2020), but did not compare the size of choice-history effects in coupled vs. uncoupled task variants in the same perceptual task.

While serial dependencies may be suboptimal in typical laboratory experiments, they might play a key role in adaptation to natural environmental statistics. Past studies found that history effects adapt to regularities in stimulus sequences (Abrahamyan et al., 2016; Braun et al., 2018; Busse et al., 2011; Hermoso-Mendizabal et al., 2020). These findings have been interpreted to reflect the learning of a prior belief about these stimulus regularities. However, the previous investigations used tasks with consistent mapping between stimulus categories and motor responses. It is thus unclear, whether this adaptation reflects learning of the regularities of the sensory input or of the actions the subject performed.

To address both these gaps, we compared choice-history effects between animals performing perceptual decision-making tasks, in which the mapping between stimulus categories and motor responses was kept constant (coupled variant) with those in which the mapping was randomized across trials (uncoupled variant). The task structure was otherwise identical between the two variants. We found that choice-history effects, and their contribution to choice behavior diminished substantially when perceptual choice was decoupled from motor response. We replicated these findings in two different tasks, performed by two different groups of animals across independent labs. Not only were these effects unexplained by differences in stimulus sequence regularities between task-variants, animals in uncoupled tasks also showed minimal adaptation to experimentally introduced stimulus sequence regularities over a period of two months. These findings suggest that the development and expression of beliefs regarding the temporal structure of the environment is influenced by priors mapped onto action-selective processes, providing further evidence for the tight link between cognitive and motor processes.

## Materials and Methods

### Animals

We conducted behavioral experiments with eight adult rhesus macaques (*Macaca Mulatta*; three females: AN, NA, BR and five males: SM, RU, DU, MA, KI). All training, surgical procedures, and experimental procedures were performed in accordance with the U.S. Public Health Service policy on the humane care and use of laboratory animals, and all protocols were approved by the National Eye Institute Animal Care and Use Committee, the University of Washington Institutional Animal Care and Use Committee, and the Columbia University Institutional Animal Care and Use Committee. Specific procedures were described in detail elsewhere (Nienborg and Cumming, 2009; Quinn et al., 2021; Shushruth et al., 2018, 2022).

### Behavioral Tasks

Throughout we refer to the categorical decision about the stimulus as the ‘choice’, and to the action reporting this choice as the ‘response’. We trained macaques on two different perceptual decision-making tasks, each with a coupled variant, in which the choice-to-response mapping was fixed, and an uncoupled variant, in which the choice-to-response mapping was varied from trial to trial (see below). Each animal performed one variant of a task.

### Disparity Discrimination

#### Coupled Disparity Discrimination Task

Two macaques (DU: number of sessions = 51, range of successful trials across sessions = [92, 3299], average number of successful trials per session = 640; RU: number of sessions = 46, range of successful trials across sessions = [96, 2294], average number of successful trials per session = 626) were trained on a coarse disparity discrimination task described previously in detail (Nienborg and Cumming, 2009). Briefly, trials initiated once the animal fixated on a central fixation point. Shortly after fixation acquisition, a dynamic circular random dot stereogram stimulus was displayed for two seconds. Animals determined whether the center of the stimulus appeared near (negative disparity) or far (positive disparity) relative to the surrounding annulus, which was displayed in the plane of the monitor. After the stimulus presentation, two dots representing choice targets appeared three degrees above and below the fixation point. Here, the upper target corresponded to a far choice, and the lower target to a near choice. The strength of the stimulus (percentage of frames with near or far disparity), and disparity of the stimulus was varied randomly across trials. The macaques received a liquid reward if they made a saccade to the correct choice target within 500ms of the target appearance. Zero-signal trials were randomly rewarded 50% of the time. We implemented a reward multiplier if animals made three consecutive correct choices by dispensing a higher reward (3x) starting with the fourth consecutive correct choice until the next error. The multiplier was reset to low (1x) following errors.

#### Uncoupled Disparity Discrimination Task

Two macaques (MA: number of sessions = 47, range of successful trials across sessions = [1490, 3408], average number of successful trials per session = 2457;, KI: number of sessions = 38, range of successful trials across sessions = [106, 1081], average number of trials per session = 623) were trained on a coarse disparity discrimination task similar to the above but with random choice-response mapping across trials, previously described in detail (Quinn et al., 2021). The task structure is similar to the coupled disparity discrimination task described above with two major differences: (i) we displayed two dynamic random dot stereograms during the stimulus presentation period: one task-relevant and one distractor. Both stimuli were visually and statistically identical on average, but their signal strengths varied independently across trials (typical set of signal strengths used in control sessions for animal KI = [0%, 5%, 10%, 20%, 30%]). For four control sessions in which we sought to lower the cognitive load for the animal during the uncoupled task, we increased the signal strengths by roughly a factor of two (set of signal strengths used in control sessions for animal KI = [2%, 10%, 20%, 40%]). We varied the location of the task-relevant and distractor stimuli between the left and right hemifields in a block-wise fashion (50 trials per block), and a change in block was indicated using “instruction trials” at the beginning of each block. During these instruction trials, we present a single task-relevant stimulus on the task-relevant hemifield for three trials. (ii) choice target icons were high signal near/far random dot stereograms, not single dots, that appeared 3-5 degrees above or below the fixation point. Animals reported their decision with a saccade to the corresponding choice target icon. Importantly, unlike the coupled variant, the location of the near/far choice targets varied randomly between the two locations from trial to trial. If animals reported the correct decision within two seconds following the onset of the targets, they received a liquid reward, similar to the coupled variant described above.

#### Stimulus Sequence Regularity Manipulation

To test if choice-history effects in the animals’ behavior reflect stimulus statistics on a particular session (Abrahamyan et al., 2016; Braun et al., 2018), we introduced stimulus regularities in individual sessions by increasing the probability of alternation of the stimulus category on subsequent trials by five percent in animals performing the uncoupled variant of the disparity discrimination task. Before each session during the manipulation period, we predefined the stimulus sign sequence in a pseudorandom fashion to introduce higher likelihoods of alternation of stimulus sign to a maximum of 55% stimulus alternation probability. We restricted further analyses of the sessions to the second half of the manipulation period when the animals had been exposed to this manipulation for at least one month and were experiencing the strongest stimulus sequence regularities of the experimental period (MA: number of sessions = 14, average number of trials per session = 1101, range of trials across sessions = [807, 1416], average signed probability of alternation of stimulus = 53.7%, range of introduced probabilities of stimulus alternation = [50.3%, 55.6%]; KI: number of sessions = 18, average number of trials per session = 679, range of trials across sessions = [297, 858], average signed probability of alternation of stimulus = 53.0%, range of introduced probabilities of stimulus alternation = [48.7%, 56.0%]). When examining the consequence of this manipulation, we compared the behavior in these sessions to the previously collected uncoupled behavioral sessions described above.

### Motion Discrimination

#### Coupled Motion Discrimination Task

One male and one female macaque (NA: number of sessions = 31, range of successful trials across sessions = [100, 1281], average number of successful trials per session = 328; BR: number of sessions = 16, range of successful trials across sessions = [110, 548], average number of successful trials per session = 193) were trained to report the direction of motion of a random dot motion stimulus, described previously in detail (Shushruth et al., 2018). Briefly, after the animals achieved fixation on a central point, two choice targets (red dots) appeared on the screen along the axis of the two potential directions of motion. After a variable delay, the random dot motion stimulus was presented on the screen. Animals were allowed to passively view the stimulus until they made a decision and reported it with a saccade to the targets in that direction. The strength of the stimulus (percentage of dots moving coherently in one direction), and direction of coherently moving dots was varied randomly across trials. Animals received liquid rewards for correct choices, and zero signal trials were rewarded at random on half of the trials. In this task, the motion direction and targets to report the choice were coupled, e.g., saccades towards an upper target corresponded to a choice reporting upward motion, and saccades towards a downward target to a choice reporting the downward direction.

#### Uncoupled Motion Discrimination Task

Two female macaques (AN: number of sessions = 28, range of successful trials across sessions = [182, 578], average number of successful trials per session = 379; SM: number of sessions = 30, range of successful trials across sessions = [165, 677], average number of successful trials per session = 388) were trained on a motion direction discrimination task, similar to the coupled task above, with two major differences: (i) the direction of coherent dots was limited to the horizontal axis (left or right) (ii) two colored choice targets were presented from a set of 3-8 possible locations on the screen. Each colored target corresponded to one motion direction, and animals made a saccade to the targets after stimulus offset. The task was described in detail elsewhere (Shushruth et al., 2022). In this task, the motion direction and targets to report the choice were uncoupled: the target locations were randomized across trials so animals could not form an action plan to report their perceptual choices during stimulus presentation.

### Analysis

#### Model-free quantification of serial dependencies

We quantified choice-history effects in a model-free manner as the horizontal separation (bias) between psychometric curves conditioned on previous choice (Wichmann and Hill, 2001). Choice-history effects are thus in units of percent signal of stimulus. While this method provides an estimate of choice-history effects, it does not factor in systematic regularities in the stimulus sequence or tease apart the relative contribution of different covariates that give rise to these biases. To this end, we used statistical modeling approaches.

### Model-based quantification of serial dependencies

#### Statistical Modeling

We fit binomial Generalized Linear Models (GLM) using different covariates to predict the probability of the animals’ choices on any given trial (Fründ et al., 2014). The covariates used in the analysis were: (i) signed stimulus strength on the current trial (S_n_), (ii) choice on the previous trial if it was rewarded (CR_n-1_), (iii) choice on the previous trial if it was an error (CE_n-1_), and (iv) reward on the previous trial (R_n-1_). We refer to the latter three regressors as history regressors. All regressors were standardized before fitting. Separate models were fit to each individual session.

Additionally, for the uncoupled task variants, we quantified the effect of motor responses by adding the following covariates: (i) spatial bias, (ii) the response on the previous trial if it was rewarded (RR_n-1_), and (iii) the response on the previous trial if it was an error (RE_n-1_). Across both uncoupled task-variants, we used these covariates in addition to the history and stimulus regressors outlined above to predict the probability of the animals’ perceptual choice on any given trial. We designed the response regressors such that for the spatial bias a positive regression coefficient indicates an upwards spatial bias, and a negative regression coefficient indicates a downward spatial bias. Conversely, for RR_n-1_ and RE_n-1_ a positive regression coefficient indicates a tendency to repeat the previous response while a negative regression coefficient reflects the tendency to alternate from the previous response. This was achieved by mapping motor responses onto the perceptual choice axis. For example, in the disparity discrimination task, this meant multiplying the previous response (up = 1, down = -1) by the perceptual choice represented by the icon in the upper hemifield on the current trial.

#### Model fitting procedure

We used nested ten-fold cross-validation to prevent overfitting. The data within each session was split into ten equally sized, continuous, and non-overlapping blocks of trials. Within each fold of cross-validation, we estimated cross-validated model parameters by fitting the model to signal trials within the training set. We used lasso regularization to perform automatic feature selection and incentivize simplicity (Tibshirani, 1996). The parameters from the training fold (90% of trials) were used to obtain behavioral predictions of choice on the leftover tenth non-overlapping block. We repeated this procedure ten times to obtain cross-validated model predictions on all trials (mean session-level variance explained for the psychometric curve across all coupled sessions = 92.5%, and for all uncoupled sessions = 91.0%). Model fitting was performed using custom-written MATLAB scripts and the built-in lassoglm function.

While this implementation does not estimate lapse rates (Fründ et al., 2014; Gupta et al., 2024), we obtained qualitatively similar results in models with lapse rates (Wichmann and Hill, 2001).

#### Choice Correlations

We quantified choice-history effects using a model-based metric we refer to as choice correlation (Lueckmann et al., 2018). We quantified the session-wise choice prediction performance of the GLM as the area under the receiver-operating characteristic curve (auROC). We converted the auROC to a choice correlation using a previously derived linear approximation (Pitkow et al., 2015). Lastly, choice correlations were transformed to Fisher z-scores. Upon conversion and transformation, a choice correlation of zero indicates an auROC value of 0.5. On zero signal trials, the choice correlation quantifies the ability of history regressors to predict the choice behavior of the animal.

Choice correlations only provide information about the choice prediction performance of the history regressors and not necessarily the type of strategy used (e.g. “win-stay” or “lose-switch”) and are thus distinct from regression coefficients whose sign indicates a tendency to repeat or switch (Abrahamyan et al., 2016; Braun et al., 2018; Fründ et al., 2014). To examine the specific strategies used by the animals, we quantified the unique contribution to the prediction from each individual regressor by assessing the variance explained by each regressor. We first computed the cross-validated choice correlation for a model fit with all regressors. We then obtained predictions from a reduced model using all regressors but one. Subtracting the choice correlation of the reduce model from the overall choice correlation yields the unique contribution by the regressor of interest. We excluded the intercept term of the generalized linear model when obtaining predictions on individual trials.

### Stimulus Regularities

We calculated stimulus alternation rate as the number of trials that the sign of the stimulus alternates from the preceding stimulus divided by the total number of opportunities for the stimulus to alternate in that session [number of alternations/(number of trials-1)]. We calculated the stimulus alternation rate for each session, subtracted 0.5, and took the absolute value to obtain the unsigned stimulus sequence regularity metric. For this transformed metric of stimulus regularity, a value of zero indicates random stimulus sequence presentation and a positive value indicates that there were stimulus sequence regularities within the session.

### Session inclusion criteria

Sessions were included in the analyses if they satisfied the following criteria. First, a session needed to contain a minimum of one hundred valid trials. Second, we excluded sessions in which the accuracy of that session was an outlier in the distribution of accuracies across sessions within an animal. Specifically, we determined the distribution of accuracies for each animal and included sessions for which accuracies fell within 4 x the interquartile range around the median. Third, the session needed to contain more than ten zero-signal trials. Lastly, the lapse rates within a given session had to be below 30%. Sixty-nine of the 417 sessions were excluded by these criteria. We obtained qualitatively similar results after relaxing the zero-signal trial and lapse rate criteria.

### Statistics and Reproducibility

The main dataset consists of 348 sessions (287 sessions for the main analyses and 61 sessions for analyses of increased stimulus regularities) across eight adult rhesus macaques (*Macaca mulatta*) grouped into four different task-variants.

We used nonparametric rank paired (Wilcoxon Signed-Rank) or unpaired (Wilcoxon Rank-Sum) tests for statistical hypothesis testing. We applied a Bonferroni correction when testing multiple hypotheses simultaneously.

## Results

In this study, we investigated whether choice-history effects—the influence of previous choices on current choices—depend on the coupling between the choice and the motor responses used to report them. To this end, we examined choice-history effects in eight macaques (*Macaca mulatta*) performing “coupled” and “uncoupled” variants of two different two-alternative forced-choice perceptual decision-making tasks (total number of successful trials across animals for coupled variant = 74,725 and for uncoupled variant = 161,376) using datasets from previously published work. Briefly, four of these animals performed two variants of a coarse disparity discrimination task (Nienborg and Cumming, 2009; Quinn et al., 2021) (**Figure 1A**), and the four other animals performed two variants of a random dot motion direction discrimination task (Shushruth et al., 2022, 2018) (**Figure 1B**). Within each task, two animals performed a “coupled” variant and two animals performed an “uncoupled” variant. In the coupled variant of the tasks, the choice-response mapping was fixed across trials and sessions. That is, each stimulus category was consistently linked to a specific motor response. Hence, the animal knew the motor response to report the choice already during the stimulus presentation (**Figure 1, left**). In the uncoupled variant, the choice-response mapping was varied randomly across trials in a session and revealed only after stimulus presentation. This required the animal to make an abstract categorical decision while the stimulus was presented and withhold motor response preparation until the targets were revealed after the stimulus presentation (**Figure 1, right**).

**Figure 1:**
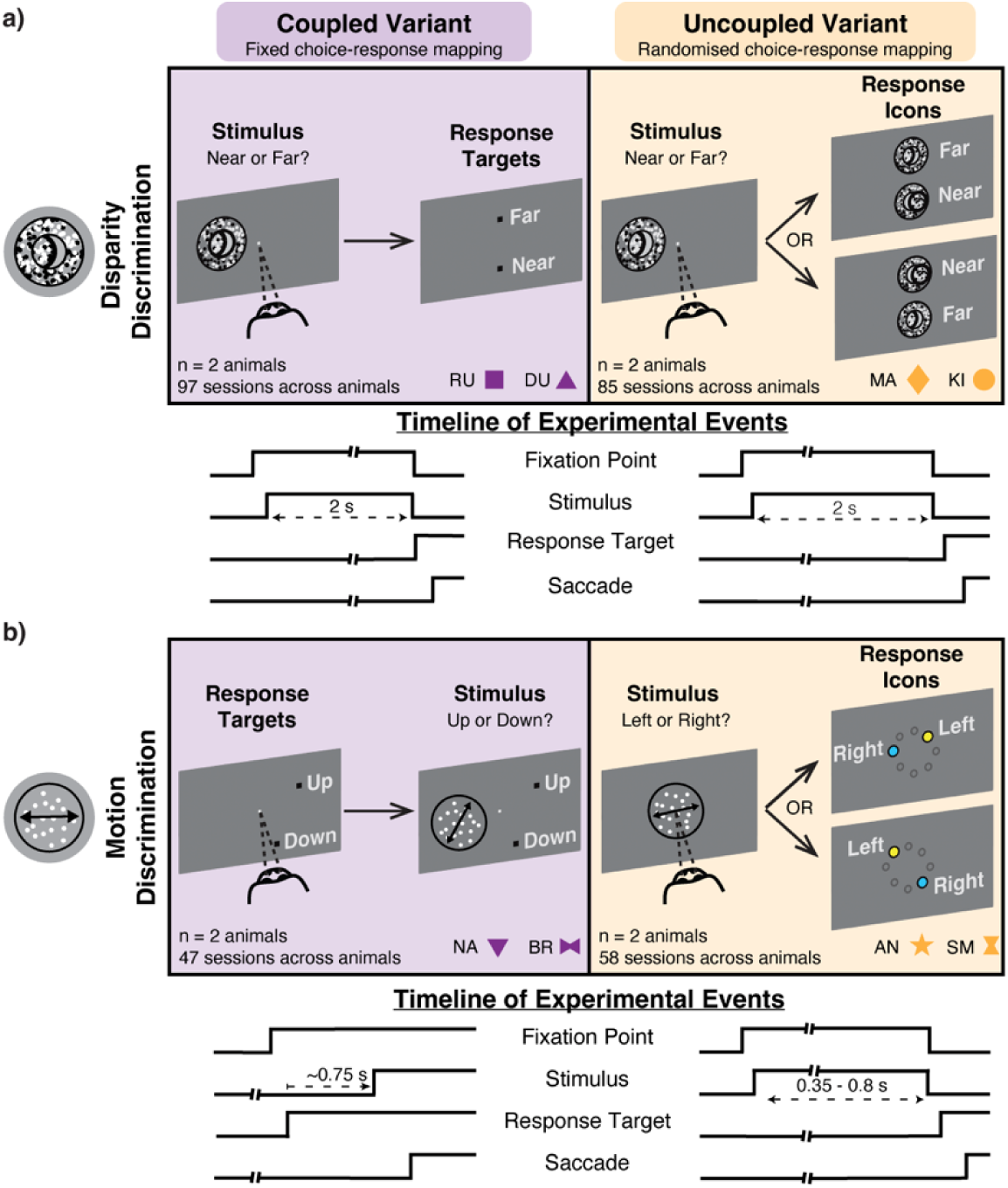
Task schematics for all datasets. A: Disparity discrimination task: In both coupled and uncoupled disparity discrimination task-variants, the animals initiated trials by acquiring fixation on a central fixation point. Upon fixation, we presented random dot stereogram stimuli for two seconds. In the coupled task-variant, animals reported a far choice with a saccade to a target in the upper hemifield and a near choice with a saccade to a target in the lower hemifield. In the uncoupled task-variant, animals reported a far choice with a saccade to the choice icon with far disparity. The location of far/near choice icons were randomly varied between two horizontally symmetric locations in the upper and lower hemifields. **B: Random dot motion direction discrimination task:** In both task-variants of the motion discrimination task, animals initiated trials by acquiring fixation on a central fixation point. In the coupled task-variant, two choice targets were then presented along an axis parallel to the two possible motion-directions in the stimulus. After a short delay, the random dot motion stimulus was presented, and animals were free to indicate their decision by making a saccade to one of the targets in a reaction-time task. In the uncoupled task-variant, the random dot motion stimulus was presented over the fixation point after a short delay following the fixation. Shortly after the stimulus offset, two icons—blue and yellow targets—appeared at two random locations sampled from a set of predetermined locations and animals responded when ready in a reaction-time task. Each direction of motion was associated with a target color, and the animals had learnt this association prior to data collection.

### Choice-history effects decreased when choice and action were decoupled

We observed choice-history biases, which decreased in uncoupled task-variants. To examine whether past choices had a systematic effect on the behavior in the current trial, we computed psychometric curves conditioned on the choice in the preceding trial. We fit psychometric curves on three different sets of trials: all trials, trials following a choice towards category #1, and trials following a choice towards category #2 (**Figure 2A**; see Materials and Methods and **Figure S1** for psychometric curves of all animals).

We found systematic shifts in the choice-history conditioned psychometric curves in the coupled variants of both tasks, consistent with prior work (Braun et al., 2018; Lueckmann et al., 2018). Conceptually, the distance between these choice-history conditioned curves represents a behavioral bias associated with the influence of previous choices on the current choice. We quantified this bias as the horizontal separation (percentage stimulus signal) between the choice-history conditioned psychometric curves (mean choice-history effect across sessions and animals performing coupled task-variants = 6.8%; range of within-animal mean choice-history effect = 2.6%, 8.6%]). But in the uncoupled tasks this analysis revealed substantially reduced magnitudes of choice-history effects (mean choice-history effect quantified as % signal shift across sessions and animals performing uncoupled task-variants = -2.2%; range of within-animal mean choice-history effect = [-2.7%, -1.6%]) (**Figure 2B**). This difference between coupled and uncoupled variants was highly significant for the disparity discrimination task (coupled vs. uncoupled: p = 8.9e-14), and followed a similar trend for the motion discrimination task (p = 0.11).

The reduced choice-history effect in uncoupled tasks variants is even more apparent in a model-based quantification. In principle, the shifts between the choice-history-conditioned psychometric functions could merely reflect regularities in the stimulus sequence rather than choice-history effects. Conversely, stimulus regularities might also obscure more pronounced choice-history effects if both shift the psychometric functions in opposite directions. To address these potential issues, we used a model-based quantification of history effects to tease apart the role of the stimulus and other covariates on behavior. We fit a generalized linear model to predict the choice behavior of animals using experimental covariates representing both current stimulus and recent choice history as regressors. We quantified the choice predictive performance of these models as the area under an ROC curve (auROC) (**Figure 3A**). We then converted the auROC values into a correlation coefficient, “choice correlation” (Pitkow et al., 2015), followed by a linearizing transform (Fisher z-transform). Expressing the choice prediction performance in this approximately linear scale facilitated interpretation and comparison of regressor contributions in subsequent analyses. Conceptually, a model using choice history information would have better choice predictive performance for sessions in which animals expressed stronger choice-history effects, resulting in higher choice correlations than a model without choice-history information. We used zero-signal trials when evaluating choice prediction performance since the stimulus on those trials is weakest and not informative about the correct choice. When quantifying choice-history effects in this model-based way, we again observed stronger choice-history effects for coupled compared to uncoupled variants of both tasks (**Figure 3B**). This difference was statistically significant for both the disparity and motion discrimination tasks (median cross-validated choice correlations for coupled vs. uncoupled in disparity discrimination task = 0.19 vs. 0.035, p-value = 1.7e-06; motion discrimination task = 0.24 vs. 0.084, p-value = 8.2e-03). Behavioral biases, including choice-history effects, are typically more pronounced for weak sensory stimuli when the sensory uncertainty is high. However, evaluating the model performance on all trials instead of zero-signal trials alone also revealed qualitatively similar trends (not shown).

**Figure 2:**
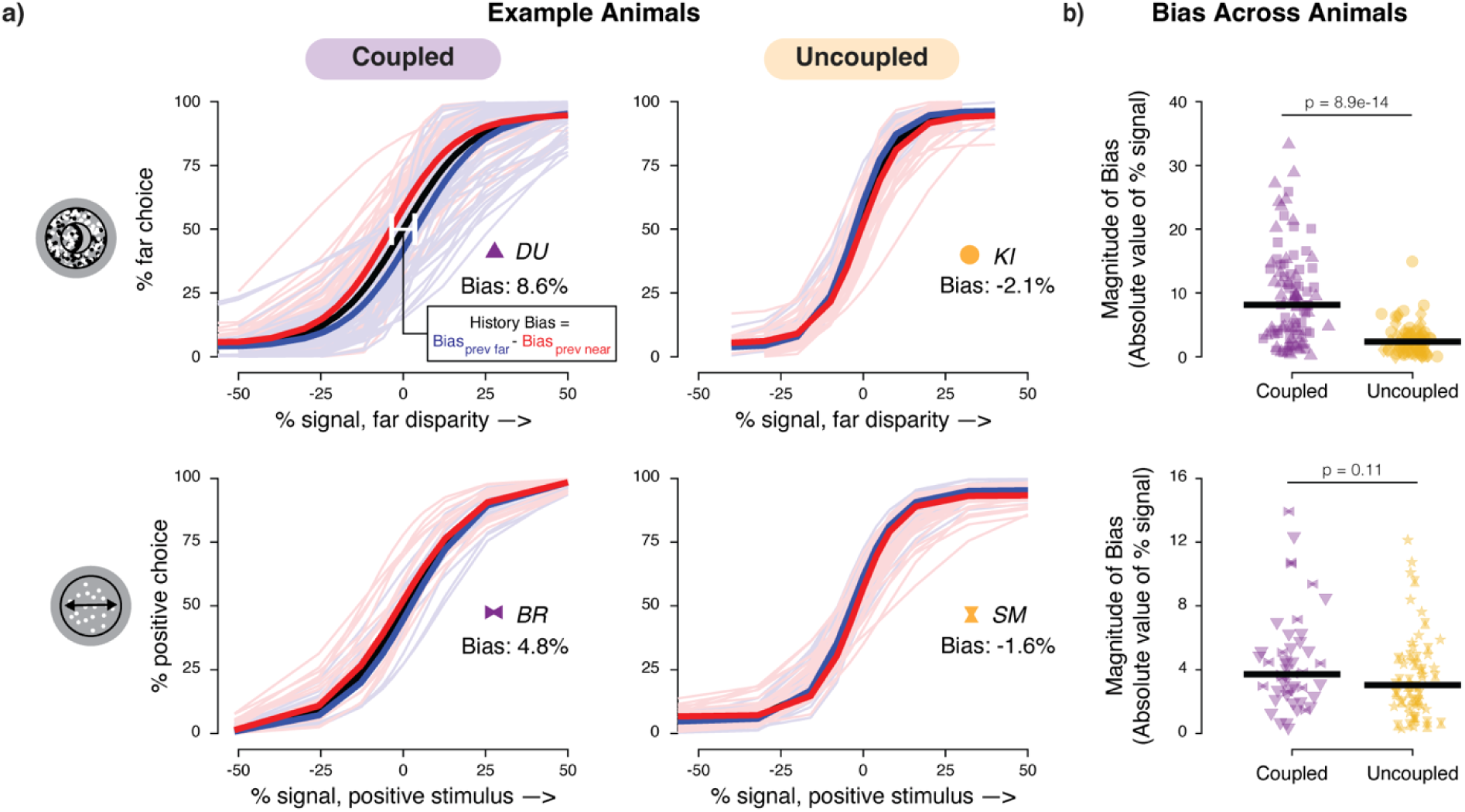
Systematic shifts in bias between choice-history conditioned psychometric curves. **A**: Example choice-history conditioned psychometric curves for four animals: two from the disparity discrimination task (top row) and two from the motion discrimination task (bottom row), with one animal each for the coupled (left column) and uncoupled (right column) variant of the task. For each session, we fit psychometric curves across three sets of trials: all trials (black), only trials where category #1 was chosen on the preceding trial (red) and only trials where the category #2 was chosen on the preceding trial (blue). Note the systematic bias (horizontal shift between the psychometric curves conditioned on previous choices) in the coupled variant, but not the uncoupled variants. See **Figure S1** for corresponding plots across all animals. Bold and thin lines depict mean across sessions for each animal and individual sessions, respectively. **B:** Magnitude of bias, quantified as the unsigned horizontal separation between choice-history conditioned psychometric curves, across all animals and task-variants. Top panel: disparity discrimination task; bottom panel: motion direction discrimination task. Data points, individual sessions.

### Action plans also do not reveal history biases in the uncoupled tasks

Since the mapping between perceptual choices and motor responses was randomized across trials in the uncoupled task-variants, we asked whether history effects were mapped to motor responses instead of perceptual choices in uncoupled task-variants. We expanded the model described above by incorporating response-history regressors and history-independent spatial bias regressors. In the uncoupled disparity discrimination task, the expanded model revealed slightly stronger choice-history effects compared to the base model. In the uncoupled motion discrimination task, no clear differences in choice-history effects were apparent between the expanded and base model (median choice correlation for models without response history regressors vs. those with response history regressors: in disparity discrimination task = 0.035 vs. 0.078, p-value = 0.017; in motion discrimination task = 0.084 vs. 0.11, p-value = 0.78). Yet, the difference in choice-history effects between coupled and uncoupled variants persisted (**Figure 3B**). Taken together, this indicates that the animals expressed neither substantial choice-history nor response-history biases when choices and the responses were uncoupled.

**Figure 3:**
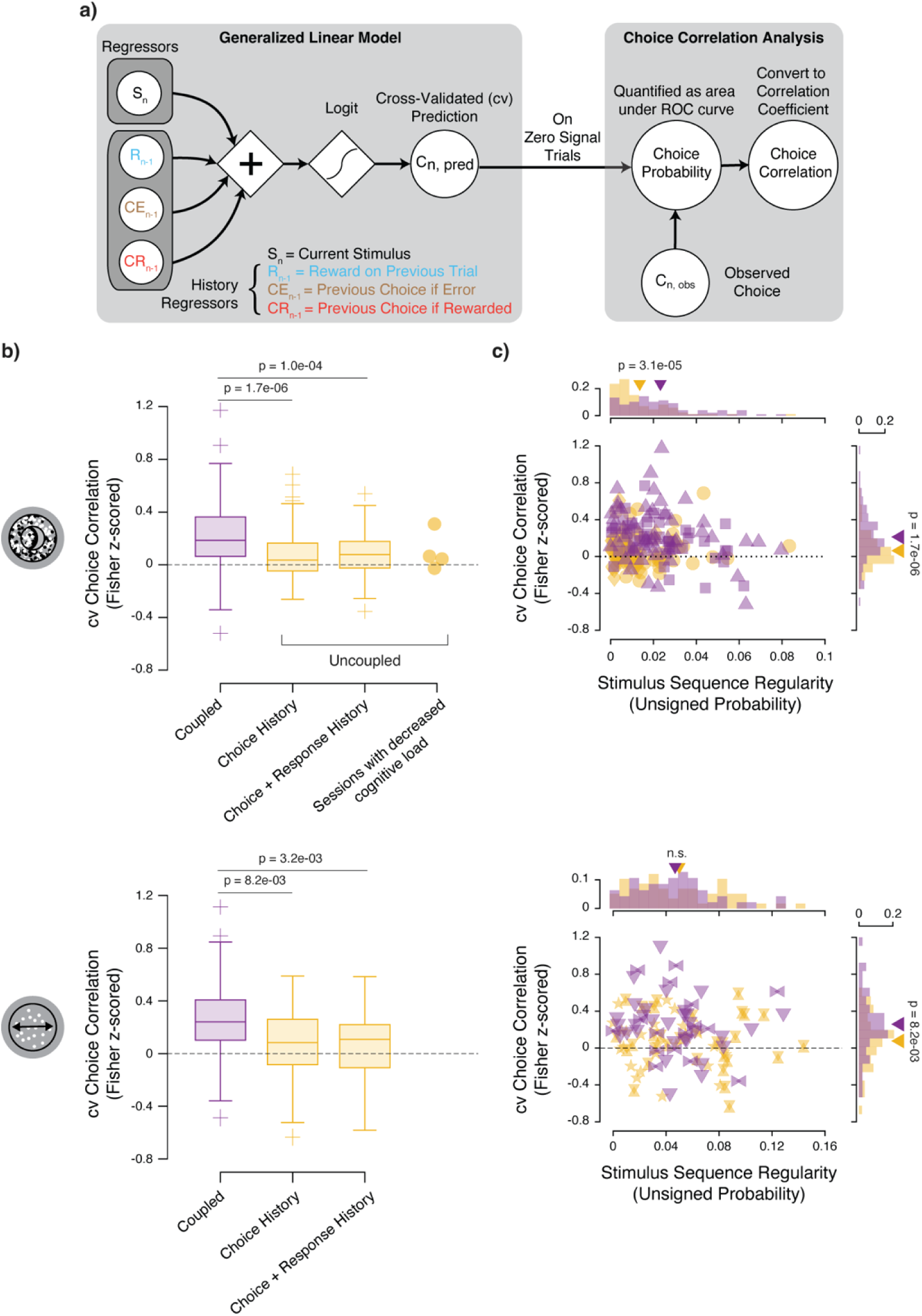
Influence of choice history on behavior is diminished substantially in uncoupled variants. **A**: Schematic of the model-based analyses to isolate the influence of choice history covariates on choice behavior. For each session, we fit a generalized linear model to animals’ choices and obtained cross-validated model predictions for each trial. We took the predictions from zero-signal trials and computed the area under the receiver operating characteristic curve, which was then transformed to choice correlation values. **B:** The distribution of choice correlation values across sessions conditioned by task-variant. Both uncoupled boxplots show cross-validated choice correlations across the same sessions but use two different models to do so: either a generalized linear model with only the current stimulus and perceptual choice history or an expanded generalized linear model with current stimulus, perceptual choice history, and motor response history. Right column shows uncoupled disparity discrimination sessions (n = 4) in which we experimentally decreased cognitive load in one animal. Top: Disparity discrimination task. Bottom: Random dot motion direction discrimination task. Box plot: whiskers, non-outlier maximum and minimum; crosses, outliers; horizontal line, median. **C:** The relationship between the unsigned stimulus sequence regularity and cross-validated choice correlation across sessions. Symbols for individual animals as in Fig. 1. Data points, individual sessions. The marginal probability histograms show the distribution of either metric. The triangles indicate the mean of the distribution. n.s., not significant.

### Differences in stimulus sequence statistics do not explain diminished choice-history effects in uncoupled tasks

The reduced choice-history effects in uncoupled tasks did also not result from task statistics. Previous studies showed that choice-history effects could reflect learned priors about environmental statistics such as stimulus sequence regularities (Abrahamyan et al., 2016; Braun et al., 2018). To confirm that the observed differences in choice-history effects between coupled and uncoupled variants was not due to differences in the stimulus statistics between the conditions, we quantified the degree of stimulus sequence regularity for each session and investigated whether choice-history effects change systematically with the regularities. We computed a stimulus sequence regularity measure—the probability of the stimulus categories to alternate/repeat more often than chance between consecutive trials— within each session. If stimulus sequence regularity influenced choice history behavior, we would expect a positive correlation between stimulus sequence regularity and choice-history effects. Contrasting with this prediction, we did not find consistent differences in stimulus sequence regularities between coupled and uncoupled variants of the tasks (95% confidence interval of bootstrapped median of unsigned stimulus regularity across sessions: in disparity discrimination, coupled variant = [0.017, 0.023], uncoupled variant = [7.2e-3, 0.012]; in motion direction discrimination, coupled variant = [0.036, 0.055], uncoupled variant = [0.032, 0.061]). Moreover, choice correlation values tended to be negatively, not positively correlated, with stimulus sequence regularities across sessions and task variants (**Figure 3C**; partial correlation between stimulus sequence regularity and choice correlation across sessions while accounting for number of trials: disparity discrimination task, coupled variant, r = -0.31, p = 1.9e-03; uncoupled variant, r = 0.036, p = 0.75; motion discrimination task, coupled variant, r = -0.14, p = 0.34; uncoupled variant, r = -0.10, p = 0.47). Together, it therefore seems unlikely that differences in task statistics between the task-variants account for the differences in choice-history effects.

### Increased cognitive load in uncoupled tasks does not explain diminished choice-history effects

Experimentally decreasing cognitive load did not reveal choice-history effects in the uncoupled variants either. It is notably more difficult for animals to be trained on and perform uncoupled tasks such as those outlined in this study (Shushruth et al., 2022) compared to coupled tasks. Animals must hold a representation of sensory evidence in mind before flexibly generating an action plan depending on the location of the presented choice targets. This suggests that the cognitive load in the uncoupled task-variants is higher than in the coupled task-variants. Prior findings observed that increased cognitive load resulted in increased choice-history effects, the opposite of our observation here (Bliss et al., 2017; Fritsche et al., 2017; Huang, 2010; Kiyonaga et al., 2017; Markov et al., 2024; Papadimitriou et al., 2016). Nonetheless, we conducted control experiments in which we experimentally reduced the cognitive load in the uncoupled disparity discrimination variant. Specifically, we modified the range of stimulus signal strengths used within a session for one animal performing the uncoupled disparity discrimination task (approximately doubling the values of signal strengths used). For the sessions with this manipulation, the animal’s choice-history effects were similar to those of the uncoupled disparity discrimination sessions (**Figure 3B**; number of sessions = 4; average number of successful trials per session = 571; mean accuracy across manipulation sessions = 86.3%; mean choice correlation computed on low signal trials discounting stimulus contributions = 0.098).

### Macaques weigh prior wins over prior losses only in coupled task-variants

The model-based quantification allowed us to pinpoint the specific, often idiosyncratic, strategies followed by the animals, and revealed that the animals tended to be primarily influenced by their previous choices when they were rewarded. We estimated the unique contribution of individual history regressors to choice behavior by computing the difference in choice correlation values of the full model with that of a reduced model fit without the regressor of interest. We found that only rewarded previous choices consistently provided a significant boost in choice correlation in the coupled variant (**Figure 4**; median unique contribution by previous choice reward in coupled task-variant: in disparity discrimination task = 0.081, p = 3.0e-10; in motion discrimination task = 0.11, p = 2.2e-03). Furthermore, animals tended to alternate choices following previously rewarded choices, as indicated by the negative coefficients for the previous choice reward regressor (**Figure S2;** mean regression coefficient for previous choice reward across sessions and animals performing coupled task-variants = -0.24; range across animals of within-animal mean regression coefficient for previous choice reward = [-0.35, -0.17]). Across both uncoupled task-variants, no choice history regressor consistently provided a boost in choice prediction performance across datasets, as expected given the near-absent choice-history effects in this condition (**Figure 4**). Furthermore, no history regressor contributed significantly to behavior in the expanded response-history model of the uncoupled datasets (**Figure S3**).

**Figure 4:**
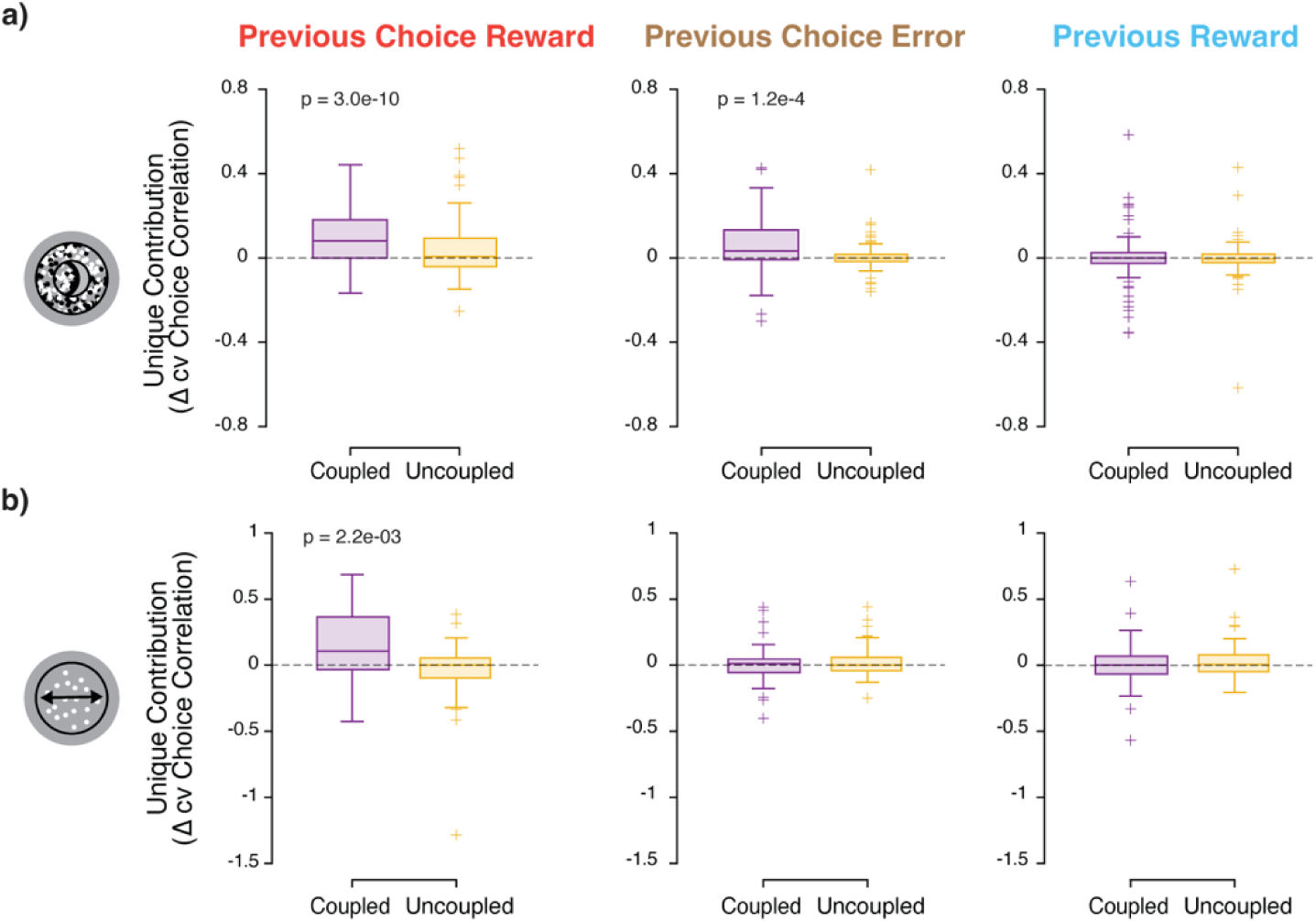
Choice-history effects in coupled task-variants are primarily explained by previously rewarded choices. Box plots show the unique contribution of each regressor to choice prediction performance for each task variant. Box plot: whiskers, non-outlier maximum and minimum; crosses, outliers; horizontal line, median. **A:** Disparity discrimination task. **B:** Random dot motion direction discrimination task. Left column of each subplot: coupled variants of the task. Right column of each subplot: uncoupled variants of the task. Bonferroni-corrected p-values compare choice prediction performance of GLM predictions with regressor of interest to choice prediction performance of GLM predictions without the regressor of interest.

### Animals showed minimal learning of stimulus sequence regularities in uncoupled tasks

Previous work in humans showed that animals adapt their dependence on choice-history to reflect experimental manipulations of stimulus sequence regularities in tasks in which choice-response mappings were fixed (Abrahamyan et al., 2016; Braun et al., 2018; Braun and Donner, 2023). Moreover, two of the animals in the coupled variant of the disparity discrimination task may have adapted to stimulus sequence regularities within those tasks (Lueckmann et al., 2018). To test if this adaptability also depends on the stable coupling between stimulus category and motor response, we introduced stronger stimulus sequence regularities across many consecutive sessions in the two animals performing the uncoupled variant of the disparity discrimination task (mean experienced stimulus alternation probability = 52.8% for MA, and 52.6% for KI; range across sessions = [50.0%, 55.6%] for MA, and [48.0%, 55.9%] for KI). Even after training the animals in the uncoupled variant in this environment for over two months (27 sessions in MA, 34 sessions in KI), neither animal expressed a significant increase in choice-history effects (**Figure 5**;mean choice correlation across sessions during second half of manipulation period vs. mean choice correlation across sessions during control period: MA = 0.059 vs. 0.069, p-value = 0.75 and KI = 0.069 vs. 0.13, p-value = 0.13). This suggests that the animals’ ability to adapt their behavior to task statistics depends on whether they are able to consistently map these statistics to potential motor responses.

**Figure 5:**
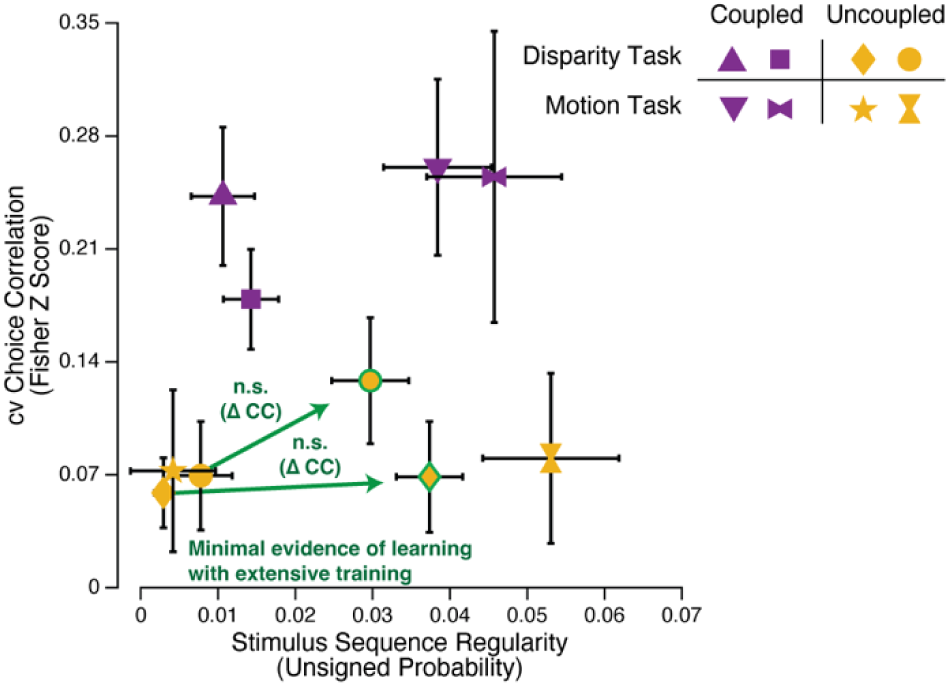
Animals express minimal evidence of learning of temporal regularities in the uncoupled condition. Each shape represents the mean stimulus sequence regularity experienced by each animal across sessions plotted against the mean choice correlation. Error bars represent standard error of the mean. We increased stimulus sequence regularities in both animals of the uncoupled variant of the disparity discrimination task (outlined in green). We only plot the second half of sessions during the two-month long manipulation period (MA = last 14 sessions of 27 manipulation sessions total, KI = last 18 sessions of 34 manipulation sessions total). Arrows connect the previously described control sessions to the manipulated sessions. Note that despite the extensive training with increased stimulus regularity, the choice-history effects remained small.

## Discussion

Here we find that choice-history effects diminish substantially when perceptual choice is uncoupled from the motor response to report it in two different perceptual decision-making tasks using non-human primates as animals. The reduction of choice-history effects in the uncoupled tasks could not be explained by several factors unrelated to the choice-response coupling. First, it did not result from reduced stimulus sequence regularities in the uncoupled tasks. Second, it was also absent when linking history effects to motor responses instead of choices. Third, the reduction in choice-history reflects the opposite pattern to that previously observed with increases in cognitive load (Bliss et al., 2017; Fritsche et al., 2017; Huang, 2010; Kiyonaga et al., 2017; Markov et al., 2024; Papadimitriou et al., 2016). The uncoupled tasks were identical to the coupled variants except for the uncoupling of choices and the motor responses to report them, implying an increased rather than lower cognitive load for the uncoupled variants. Moreover, our finding was robust in our control experiments lowering the cognitive load also for the uncoupled variant of the disparity discrimination task. Finally, the minimal choice-history effects persisted even after extended exposure to experimentally induced regularities in the stimulus sequence. Together, our findings highlight the link between perceptual choices and actions for behaviors that are often interpreted in purely cognitive terms (Gardner, 2019) or as sensory adaptations to natural environmental statistics (e.g. (Fischer and Whitney, 2014).

Our results extend findings of previous studies in humans that also used uncoupled task variants to probe history effects (Braun et al., 2018; Pape and Siegel, 2016; Zhang and Alais, 2020), by comparing the size of the effects on behavior. Previous studies found evidence for choice-history or response-history effects in behavior, but the size was not compared between the coupled and uncoupled variants, and the observed effects differed between studies. Indeed, previous studies only examined effects of previous motor responses on those in the current trial (Pape and Siegel, 2016), observed no effect of previous motor responses (Braun et al., 2018), or found opposing effects of previous motor responses to choice-history effects (Zhang and Alais, 2020). Moreover, effects were typically compared directly via the weights assigned to regressors in model-based quantifications (Braun et al., 2018; Zhang and Alais, 2020). In contrast, our study quantified the contribution of history regressors not as regression weights but as the ability of the regression model to predict choice behavior when the visual information was weakest on zero signal trials and the animals putatively relied most on their heuristics to make choices. This analysis revealed that, even when previous choice weights were non-zero (**Figure S3**) in the uncoupled task variant, history regressors still had minimal choice predictive performance.

History-based heuristics are often thought to reflect priors or learned statistical regularities of natural environments (Gardner, 2019). But the degree to which these history-based strategies depend on the stable choice-response mappings typically present in our natural environment (Cisek and Kalaska, 2010) has remained unclear. Previous work in humans identified a significant involvement of motor areas in predicting current response from previous response (Pape and Siegel, 2016) in an uncoupled perceptual decision paradigm. In coupled tasks, previous work identified sensorimotor circuits in the posterior parietal cortex that maintained a short-term memory of past choices in mice (Morcos and Harvey, 2016) and humans (Urai and Donner, 2022), and mediated history effects on behavior in rats (Akrami et al., 2018). The role of parietal cortex in representing the evolution of decisions for particular actions in coupled task is well established (De Lafuente et al., 2015; Shadlen et al., 2008; Tosoni et al., 2008). However, this representation requires knowledge of the specific motor response to report a decision and is absent for uncoupled (abstract) decisions (Shushruth et al., 2022). Our results here further emphasize the coupling of perceptual decisions to action plans, even when the decisions can be done independently of the particular action to report them. Of course, the animals need to infer the task rules during training from their actions such that their training history may emphasize the coupling of decisions and action plans. Together, these collective findings are compatible with distributed sensorimotor networks for decision-formation and other cognitive functions (Bondy et al., 2024; Findling et al., 2025; Hunt and Hayden, 2017; Okazawa and Kiani, 2023; Pinto et al., 2019; Rosen and Freedman, 2026; Ruff et al., 2025; Van Den Brink et al., 2023). In this view seemingly abstract variables such as statistical priors, past choices, or evolving decisions are not represented by specialized domain-general circuits but rather within a distributed sensorimotor network that changes depending on specific task demands and contexts.

## Acknowledgments

We thank the members of the Laboratory of Sensorimotor Research for fruitful discussions. We acknowledge funding from the National Eye Institute Intramural Research Program at the National Institutes of Health (NIH, grant no. 1ZIAEY000570-01 to H.N.), the Center on Compulsive Behaviors via the NIH Shared Resource Subcommittee (NIH, to B.C.T.), the Howard Hughes Medical Institute, and the NIH extramural program (grants: R01NS113113, R21AG067108, R01MH122513, to M.N.S.). The funders had no role in study design, data collection and analysis, decision to publish or preparation of the manuscript. This research was supported by the Intramural Research Program of the National Institutes of Health (NIH), The National Eye Institute. The contributions of the NIH author(s) were made as part of their official duties as NIH federal employees, are in compliance with agency policy requirements, and are considered Works of the United States Government. However, the findings and conclusions presented in this paper are those of the author(s) and do not necessarily reflect the views of the NIH or the U.S. Department of Health and Human Services.

## Contributions

B.C.T. and H.N. conceived the project and provided the methodology. S.S. and H.N. provided the data. S.T. and B.C.T carried out the investigations. S.T. analyzed the data and visualized the results. S.T., B.C.T. and H.N. wrote the original draft of the paper. S.T., B.C.T., S.S., M.N.S., and H.N. reviewed and edited the paper. H.N. and M.N.S. acquired funding and administered the project.

## Supplementary Figures

**Supplementary Figure 1:**
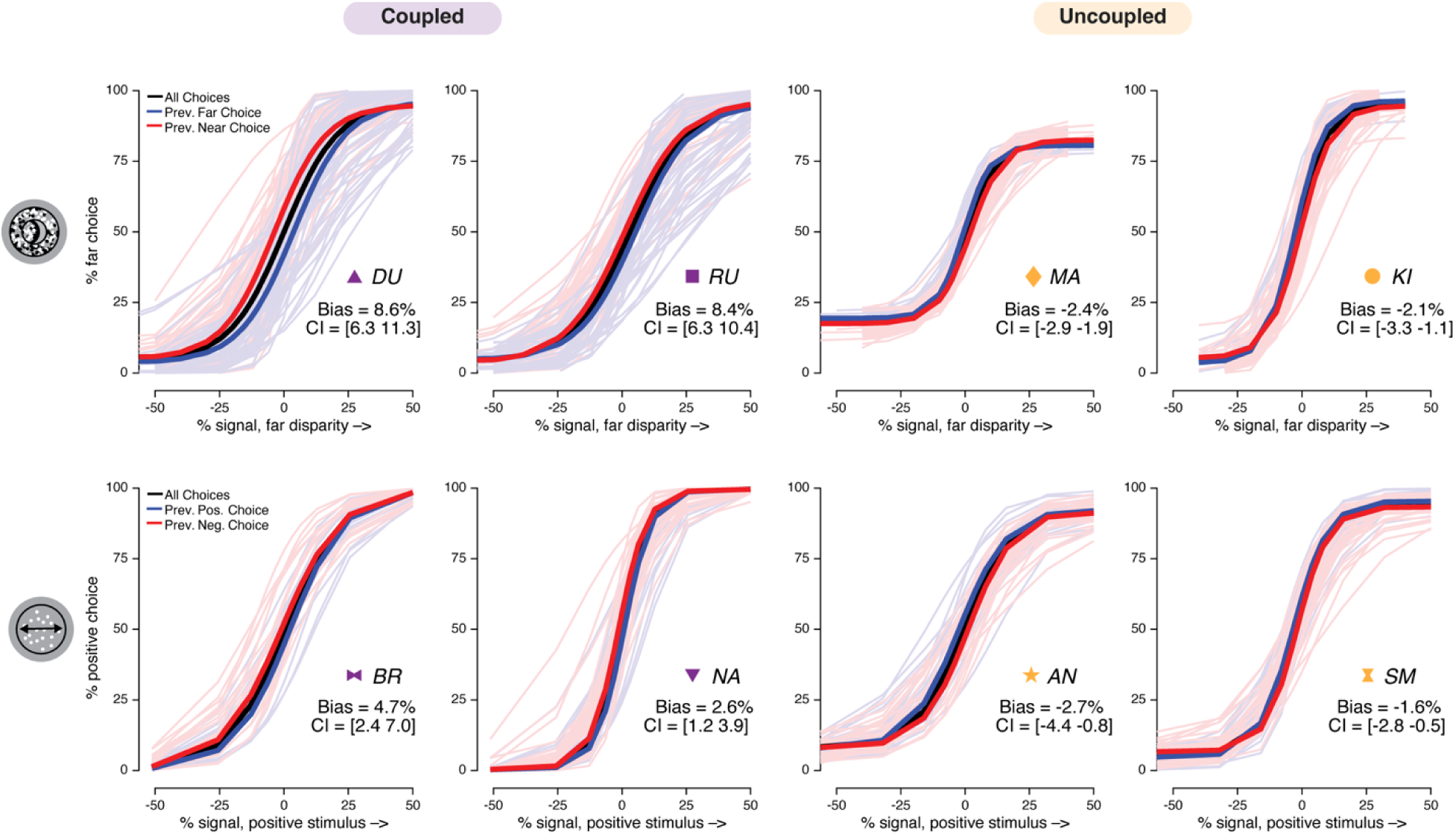
Choice-History Conditioned Psychometric Curves for individual animals. Here, we plotted choice-history conditioned psychometric curves for all four animal performing the disparity discrimination task (top row) and the motion discrimination task (bottom row). Each animal is denoted by a two-letter abbreviation and shape-icon. Transparent traces represent individual session choice-history conditioned fits. Thick trace represents the mean fit across sessions. Please see Figure 1 and **Materials and Methods** for details on icon-animal mapping, model-fitting, sign-convention, and quantification of choice-history effects. “Bias” indicates mean choice-history effects across sessions and “CI” indicates the 95% bootstrap confidence interval for the mean bias.

**Supplementary Figure 2:**
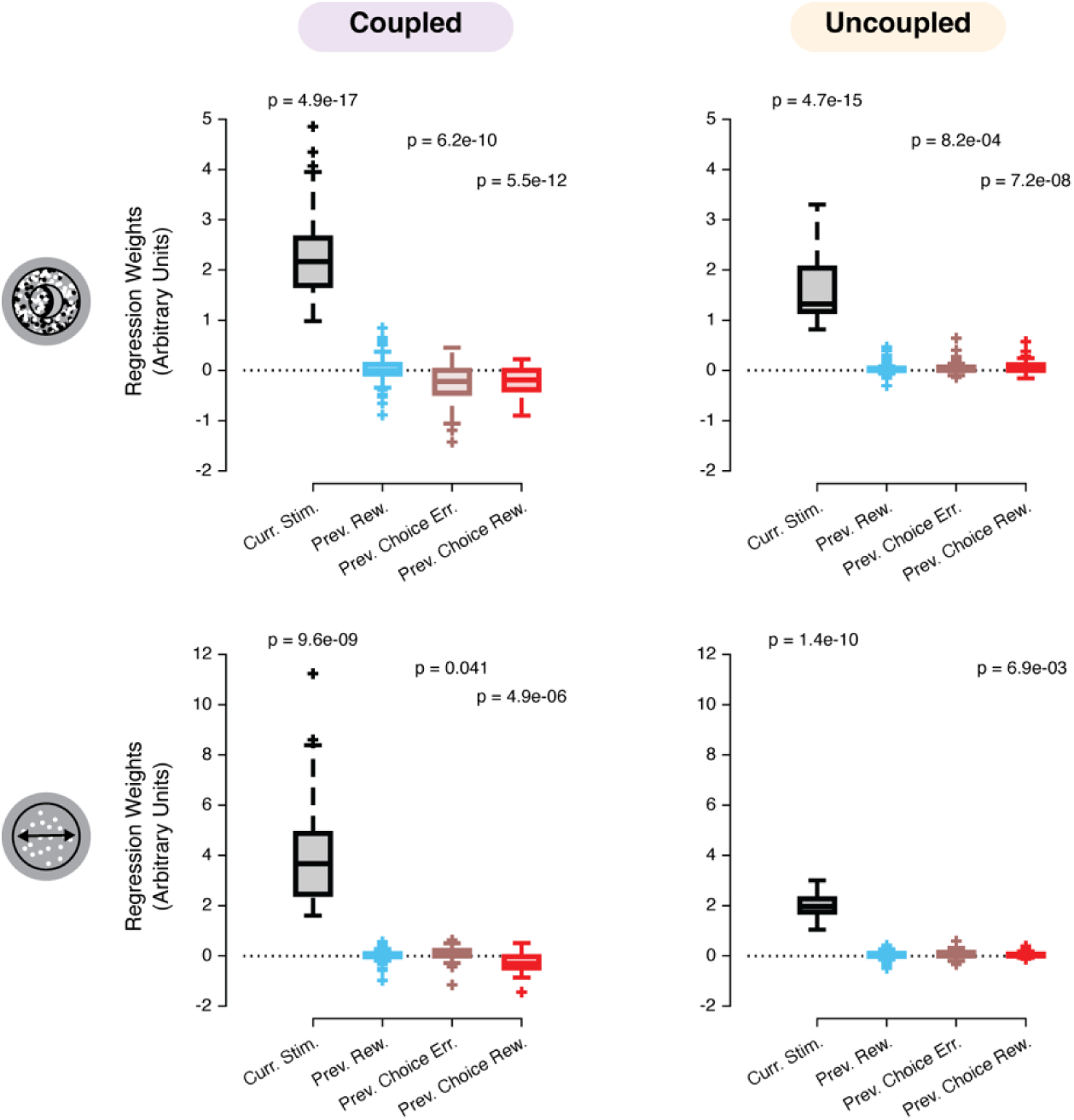
Model weights describing choice history are larger in coupled animals. We fit a cross-validated generalized-linear model using regressors representing stimulus information and choice history. We obtained regression coefficients for all regressors within each session. Here, we plotted the regression coefficients for the stimulus and history parameters across sessions for each task variant (top row, disparity discrimination task; bottom row, motion discrimination task). History regressors, whose weights differ significantly from zero do not necessarily provide a significant boost in choice correlation (compare Fig. 3). History regressors that significantly increase choice correlation had negative regression coefficient weights. It indicates that the animals tend to alternate their choices. Boxplots show the distribution of model regression coefficients across sessions conditioned by task-variant. Box plot: whiskers, non-outlier maximum and minimum; crosses, outliers; horizontal line, median.

**Supplementary Figure 3:**
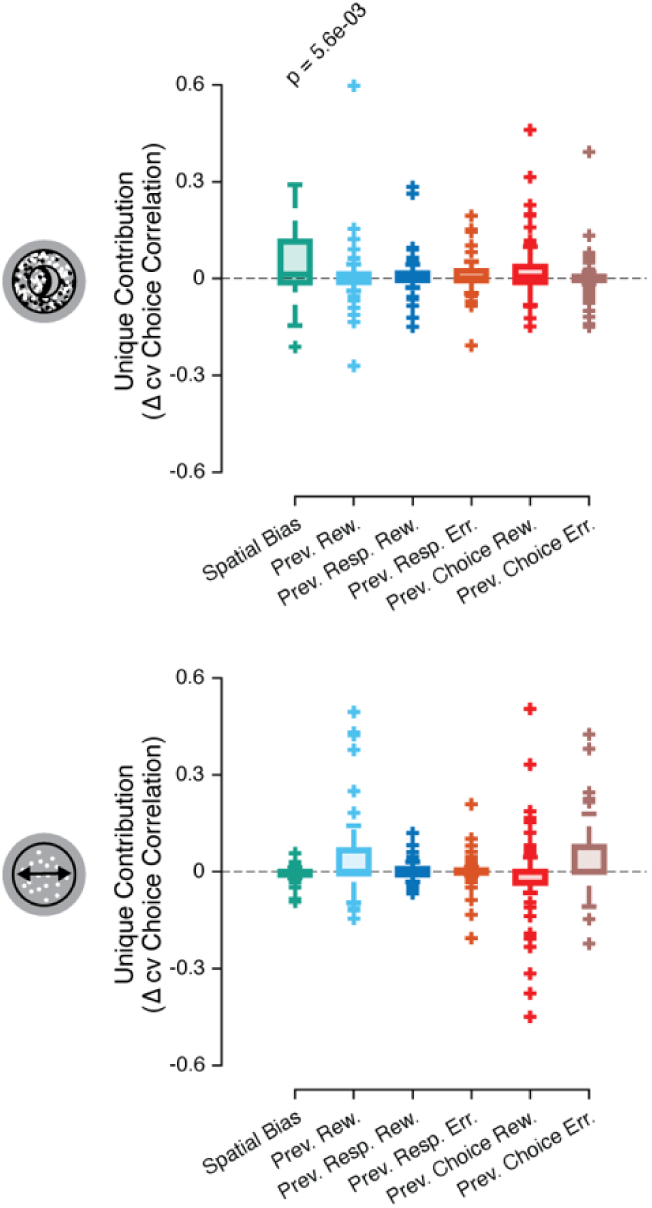
Previous motor responses and spatial biases do not account for differences in history effects between task-variants. We quantify the unique contribution to choice prediction performance of each regressor in the expanded choice and response history model (see **Materials and Methods**). Box plot: whiskers, non-outlier maximum and minimum; crosses, outliers; horizontal line, median. Bonferroni-corrected p-values represent the significance of contribution to choice prediction performance by that particular regressor (Wilcoxon Signed-Rank test).

